# Antibiotic resistance and metabolic profiles as functional biomarkers that accurately predict the geographic origin of city metagenomics samples

**DOI:** 10.1101/476853

**Authors:** Carlos S. Casimiro-Soriguer, Carlos Loucera, Javier Perez Florido, Daniel López-López, Joaquin Dopazo

**Affiliations:** Clinical Bioinformatics Area. Fundación Progreso y Salud (FPS). CDCA, Hospital Virgen del Rocio. 41013. Sevilla. Spain; Functional Genomics Node (INB). FPS. Hospital Virgen del Rocio. 41013 Sevilla. Spain; Bioinformatics in Rare Diseases (BiER). Centro de Investigación Biomédica en Red de Enfermedades Raras (CIBERER). FPS. Hospital Virgen del Rocio. 41013. Sevilla. Spain

**Keywords:** Machine learning, classification, metagenomics, whole genome sequencing, functional profiling, antibiotic resistance

## Abstract

**Background:** The availability of hundreds of city microbiome profiles allows the development of increasingly accurate predictors of the origin of a sample based on its microbiota composition. Typical microbiome studies involve the analysis of bacterial abundance profiles.

**Results:** Here we use a transformation of the conventional bacterial strain or gene abundance profiles to functional profiles that account for bacterial metabolism and other cell functionalities. These profiles are used as features for city classification in a machine learning algorithm that allows the extraction of the most relevant features for the classification.

**Conclusions:** We demonstrate here that the use of functional profiles not only predict accurately the most likely origin of a sample but also to provide an interesting functional point of view of the biogeography of the microbiota. Interestingly, we show how cities can be classified based on the observed profile of antibiotic resistances.

## Background

In recent years there has been an increasing interest in microbiome research, especially in the context of human health [1]. However, bacteria are ubiquitous and microbiotas from many different sources have been object of scrutiny [2]. Specifically, environmental metagenomics recently is gaining much attention [3]. The Metagenomics and Metadesign of the Subways and Urban Biomes (MetaSUB) is an International Consortium with a wide range of aims, currently involved in the detection, measurement, and design of metagenomics within urban environments [4]. Typically, microbiomes have been studied by analyzing microbial abundance profiles obtained either from 16S RNAs or from whole genome sequencing (WGS), which can be further related to specific conditions [5, 6]. More recently, 16sRNA data has been used as a proxy to derive functional profiles by assigning to each sample the functional properties (pathways, resistance or virulence genes, etc.) of the genomes of reference of each species identified in it [7, 8]. However, 16sRNA data does not allow direct inference of genes actually present in the bacterial population studied [9]. Contrarily, metagenomics shotgun sequencing allows inferring a quite accurate representation of the real gene composition in the bacterial pool of each sample that can be used to identify strain-specific genomic traits [10, 11]. For example, the focused study of specific traits such as antibiotic resistance or virulence genes has been used to detect pathogenic species among commensal strains of *E. coli* [12]. Also, general descriptive functional profile landscapes have been used to understand the contribution of microbiota to human disease [13]. However, in spite of the abundance of different types of metagenomics profiles in human health [12, 14], little is known on the value of existing profiling tools when applied to urban metagenomes [15].

Here, we propose a machine learning innovative approach in which of functional profiles of microbiota samples obtained from shotgun sequencing are used as features for predicting geographic origin. Moreover, in the prediction schema proposed, a feature relevance method allows extracting the most important functional features that account for the classification. Thus, any sample is described as a collection of functional modules (e.g. KEGG pathways, resistance genes, etc.) contributed by the different bacterial species present in it, which account for potential metabolic and other functional activities that the bacterial population, as a whole, can perform. We show that the functional profiles, obtained from the individual contribution of each bacterial strain in the sample, not only display a high level of predictive power to detect the city of origin of a sample but also provide an interesting functional perspective of the city analyzed. Interestingly, relevant features, such as antibiotic resistances, can accurately predict the origin of samples and are compatible with epidemiological and genetic observations.

## Material and methods

### Data

Sequence data were downloaded from the CAMDA web page (http://camda2018.bioinf.jku.at/doku.php/contest_dataset#metasub_forensics_challenge). There are four datasets: *training dataset* composed of 311 samples from eight cities (Auckland, Hamilton, New York, Ofa, Porto, Sacramento, Santiago and Tokyo, *test dataset 1*, containing 30 samples from New York, Ofa, Porto and Santiago; *test dataset 2* containing 30 samples from three new cities (Ilorin, Boston and Lisbon) and *test dataset 3* containing 16 samples from Ilorin, Boston and Bogota

### Sequence data processing

Local functional profiles were generated from the original sequencing reads by the application MOCAT2 [16] which uses several applications for the different steps. FastX toolkit is used for trimming the reads and SolexaQA [17] to keep the reads in which all quality scores are above 20 and with a minimum length of 45. In order to remove possible contamination with human genomes we screened the reads against hg19. In this step MOCAT2 use SOAPaligner v2.21 [18]. High quality reads were assembled with SOAPdenovo v1.05/v1.06 [18]. Then, genes were detected inside contigs using Prodigal [19]. Figure 1A outlines the procedure followed.

**Figure 1.**
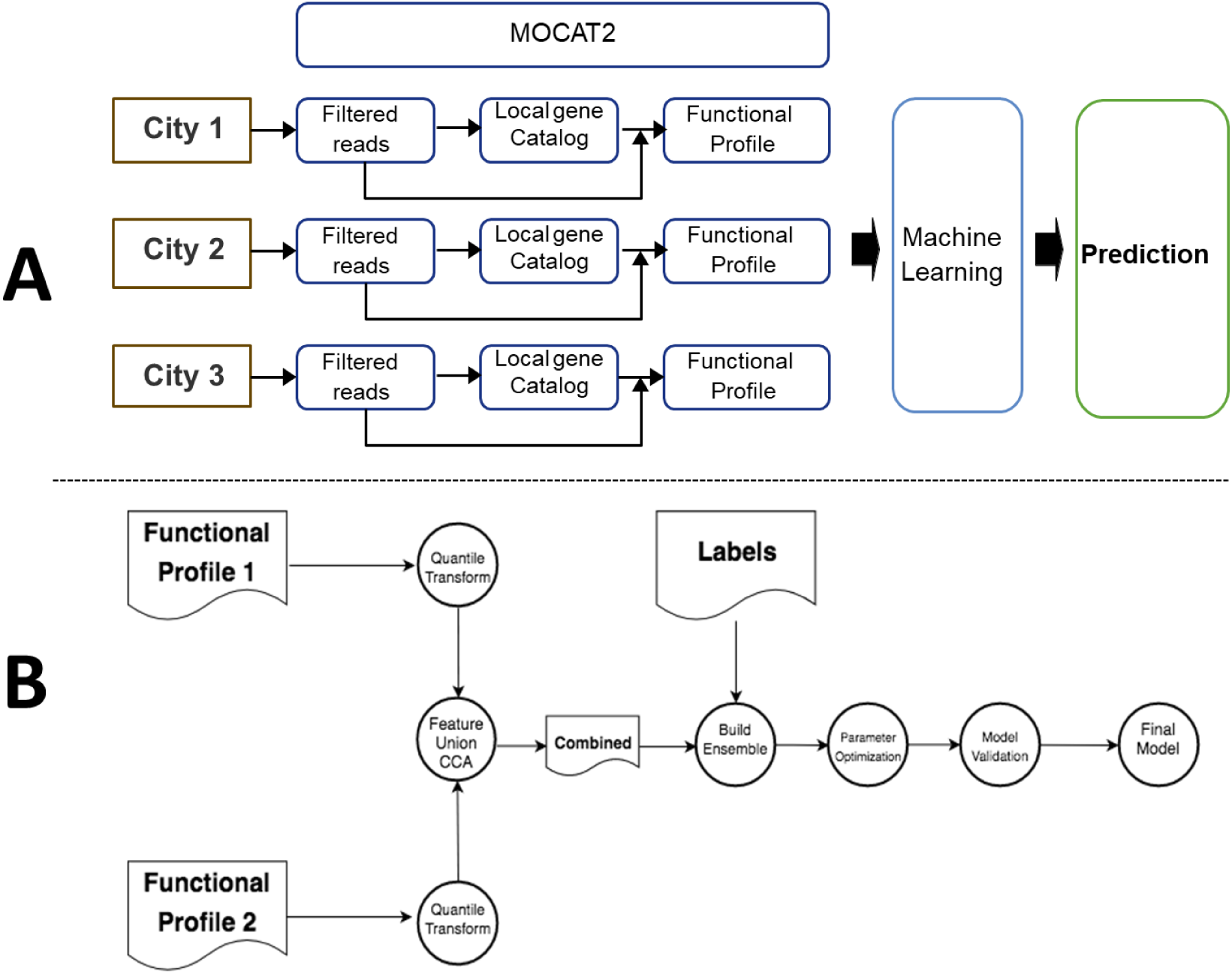
Schemas of: A) the annotation and machine learning procedure and B) the fusion pipeline, as explained in Methods.

### Functional profiles

CD-hit [20] with a 95% identity and a of 90 % overlap with the sorter sequence was used to create a local gene catalog for each city. Gene catalogs were annotated using DIAMOND (v0.7.9.58) [21] to align the genes against the orthologues groups of the database eggNOG (v4.5) [22]. MOCAT2 pre-computed eggNOG orthologous groups sequences with annotations from other databases. Thus, a functional profile is generated for each sample by assessing the gene coverage for KEGG (v74/57) [23] and CARD (August 2015) [24] functional modules. Finally each sample is normalized by the number of mapped reads against local gene catalog.

### Machine learning pipeline

The machine learning phase takes the complete KEGG Module functional profile as the input feature space, i.e. each training/validation sample is represented as a 1D-array where the values/features are a one to one map with the KEGG modules. The machine learning pipeline has been implemented in python 3.6 by making use of scikit-learn [25]. The training and validation datasets are transformed according to a quantile transformation whose parameters are learned from the training data. Subsequently, we apply the learned data representation to each validation dataset. The quantile preprocessing performs a feature-wise non-linear transformation which consists on transforming each variable to follow a normal distribution. This is a robust preprocessing scheme since the impact of the outliers is minimized by spreading the most frequent values.

In order to visualize such a high dimensional dataset we use the t-distributed Stochastic Neighbor Embedding (t-SNE) [26] methodology. Due to the fact that the feature space dimension is much greater than the number of samples, a principal component analysis (PCA) is performed to reduce the dimensionality of the embedding process carried out by t-SNE.

### Classification pipeline

To classify each sample into one of the known cities a classification pipeline was developed which mainly consists of: i) A base learner with decision trees, ii) An ensemble of base learners via Scalable Tree Boosting [27] and, iii) A Bayesian optimization framework for tuning the hyper parameters. The optimization tuning has been done by following the guidelines provided in [28].

In order to estimate the generalization error of the underlying model and its hyper-parameter search we have used a nested/non-nested cross-validation scheme. On the one hand, the non-nested loop is used to learn an optimized set of hyper-parameters, on the other hand, the nested loop is used to estimate the generalization error by averaging test set scores over several dataset splits. The scoring metric is the accuracy and the hyper-parameter learning is done on the inner/nested cross validation by means of Bayesian optimization. Figure 1A contains a schema of the whole pipeline followed here.

### Fusion pipeline

In order to improve the classification accuracy of the proposed method we can fuse different functional profiles by learning an approximation of the latent space by means of Canonical Correlation Analysis (CCA) and then applying the machine learning pipeline already proposed. Thus, a multi view classification problem, where the views are the functional profiles can be constructed. A quantile transformation is learned for each dataset as previously described (Figure 1A) and then, the latent space between both views is built by making use of CCA as previously described [29]. Finally, we apply the proposed classification pipeline (except the quantile transformation).

Given two datasets X_1_ and X_2_ that describe the same response Y, CCA-based feature fusion consists in concatenating, or adding, the latent representations of both views in order to build a single dataset that captures the most relevant patterns. CCA finds one transformation (T_i_) for each view in such a way that the linear correlation between their projections is maximized in a latent space with less features that either X_1_ or X_2_. Figure 1B shows a diagram that summarizes the Fusion Pipeline.

## Results and discussion

### The CAMDA challenge

The CAMDA challenge *test dataset* consists of 311 samples from eight cities: Auckland, Hamilton, New York, Ofa, Porto, Sacramento, Santiago and Tokyo. The predictor was trained with this test dataset and then used to predict new samples

### Classification of the cities

The sequences from the CAMDA *test dataset* were processed as described in methods and a KEGG-based functional profile was obtained for all the samples of the training datasets. The cities display characteristic functional profiles (see Figure 2) that clearly differentiate them. Figure 3 shows how the functional profiles separate the different cities as result of the application of the clustering pipeline on the *training dataset 1*. The results reveal the strong performance of the suggested pipeline as most of the classes (i.e. cities) are well separated, except for Hamilton and Auckland (both New Zealand cities) which are separated from the other cities but are very difficult to distinguish between themselves. This functional similarity was expected due to their geographical closeness and its connection. Table 1 shows the cross validation results, where the New Zealand cities could not be properly resolved as some of the samples were missassigned.

**Table 1.** Cross validation of the CAMDA training dataset.

| Truth / Pred | Auckland | Hamilton | NY | Ofa | Porto | Sacramento | Santiago | Tokyo | All |
| --- | --- | --- | --- | --- | --- | --- | --- | --- | --- |
| Auckland | 9 | 4 | 0 | 1 | 0 | 1 | 0 | 0 | 15 |
| Hamilton | 3 | 11 | 2 | 0 | 0 | 0 | 0 | 0 | 16 |
| NY | 1 | 0 | 110 | 1 | 0 | 6 | 2 | 6 | 126 |
| Ofa | 0 | 0 | 3 | 17 | 0 | 0 | 0 | 0 | 20 |
| Porto | 0 | 0 | 0 | 0 | 60 | 0 | 0 | 0 | 60 |
| Sacramento | 0 | 0 | 0 | 0 | 0 | 34 | 0 | 0 | 34 |
| Santiago | 0 | 0 | 1 | 0 | 0 | 0 | 17 | 2 | 20 |
| Tokyo | 0 | 0 | 0 | 0 | 0 | 0 | 0 | 20 | 20 |
| All | 13 | 15 | 116 | 19 | 60 | 41 | 19 | 28 | 311 |

**Figure 2.**
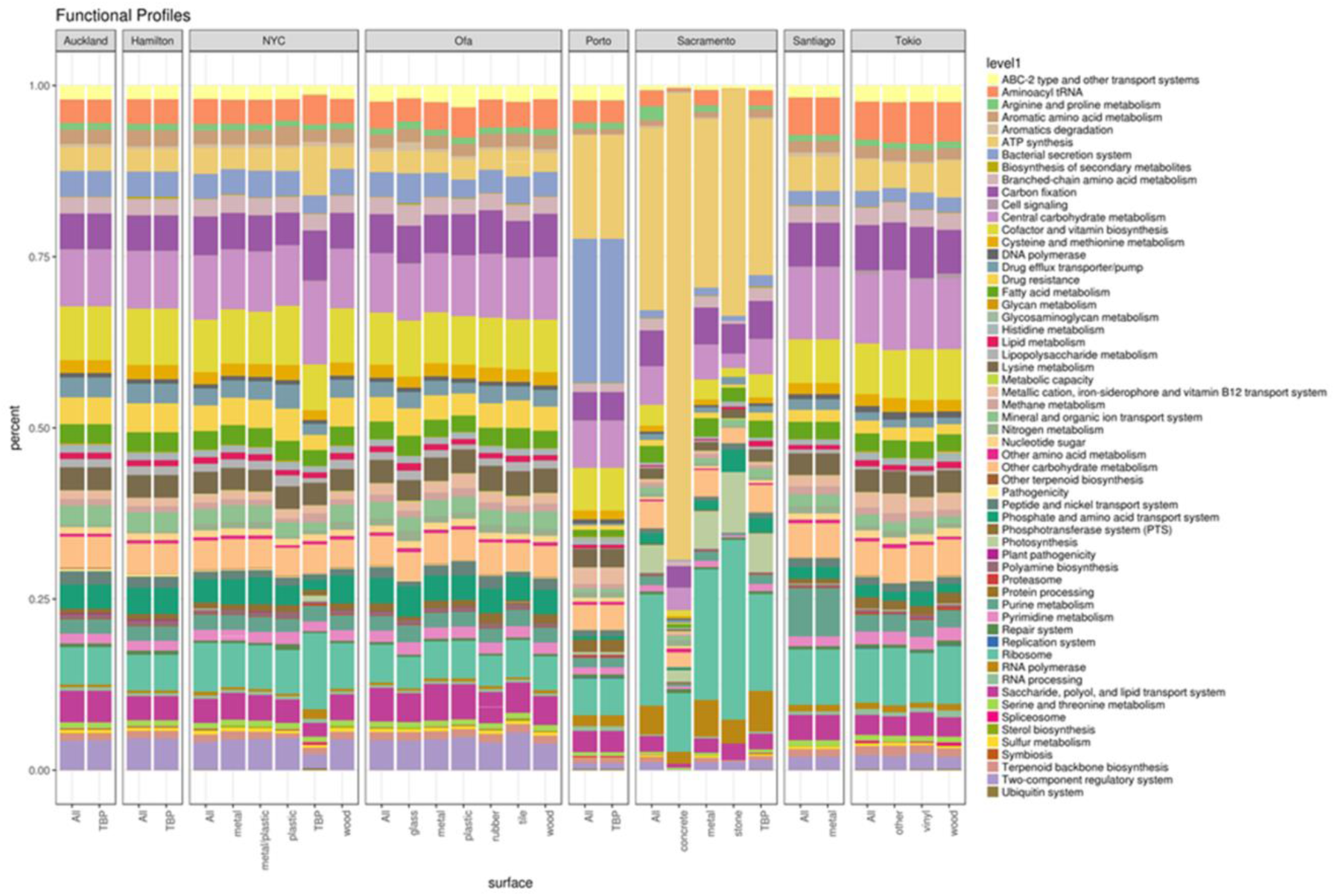
Percentages of 59 high level KEGG modules defining the functional profiles for each city and surface by city are shown (for the sake of the visualization KEGG modules were collapsed to the corresponding highest level definitions)

**Figure 3.**
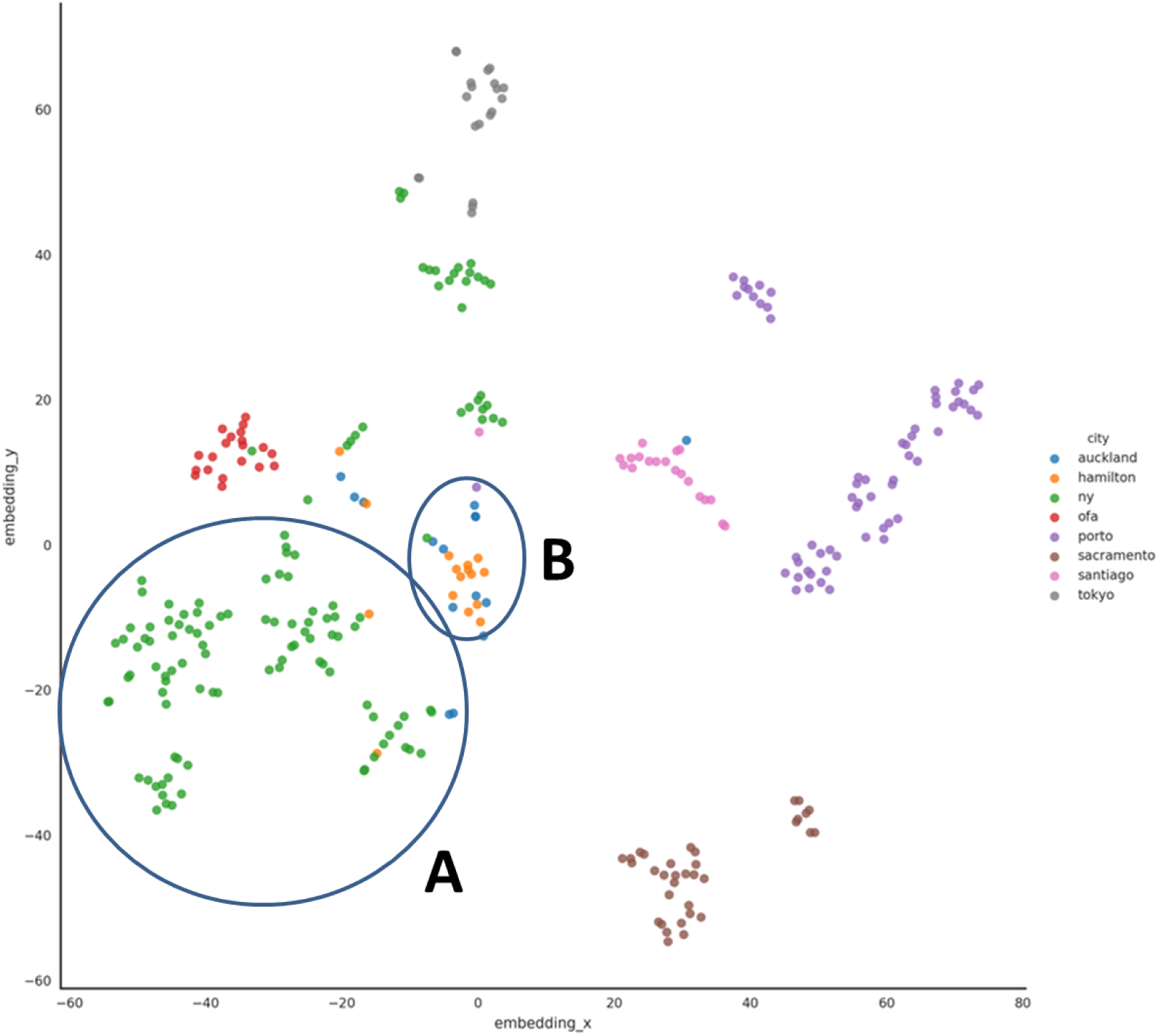
Classification of the cities of the training set based on KEGG-based functional profiles. A) As expected, the New York cluster shows the highest dispersion. B) Hamilton and Auckland (both New Zealand cities connected by a train) are separated from the other cities but are very difficult to distinguish among them.

### Feature extraction and biological relevance in the classification

An advantage of using functional modules as classification features is that their biological interpretation is straightforward. Here, the most relevant features were extracted from the classification pipeline from each run of the experiment by averaging the feature importance of each base learner of the ensemble (an easily computable scores since we use decision trees). The features that appeared in all the experiments were selected. Then, to assure the relevance of each extracted feature we cross-reference it with those found by an l1-driven logistic regression model. Finally we perform a 10-fold cross-validated prediction in order to assert that the difference in accuracy is close to that found with the whole dataset. The total number of extracted features adds up to 44.

Importantly, the features used for the classification have a direct biological meaning and account for city-specific functional properties of the bacterial samples found in each city. As an example of easy interpretation is the city of Ofa. Out of the seven features that distinguish this city from the rest of cities (see Figure 4), three KEGG modules are related with antibiotic resistances (see Table 2). Interestingly, antibiotic resistance had already been studied in the MetSUB dataset by directly searching the presence in *P. stutzeri mexA* strains (that carry the *mexA* gene, a component of the MexAB-OprM efflux system, that confer resistance to antibiotics [30]) present in samples from some cities [15]. However, in the approach presented here, that allowed the detection of the most relevant functional features that characterize cities, antibiotic resistance arises as a highly discriminative feature for some of them.

**Table 2:** The most relevant KEGG modules in Ofa

| KEGG ID | KEGG name |
| --- | --- |
| M00090 | Phosphatidylcholine (PC) biosynthesis, choline => PC |
| M00092 | Phosphatidylethanolamine (PE) biosynthesis, ethanolamine => PE |
| M00224 | Fluoroquinolone transport system |
| M00309 | Non-phosphorylative Entner-Doudoroff pathway, gluconate/galactonate => glycerate |
| M00480 | VraS-VraR (cell-wall peptidoglycan synthesis) two-component regulatory system |
| M00494 | NatK-NatR (sodium extrusion) two-component regulatory system |
| M00658 | VanS-VanR (actinomycete type vancomycin resistance) two-component regulatory system |

**Figure 4.**
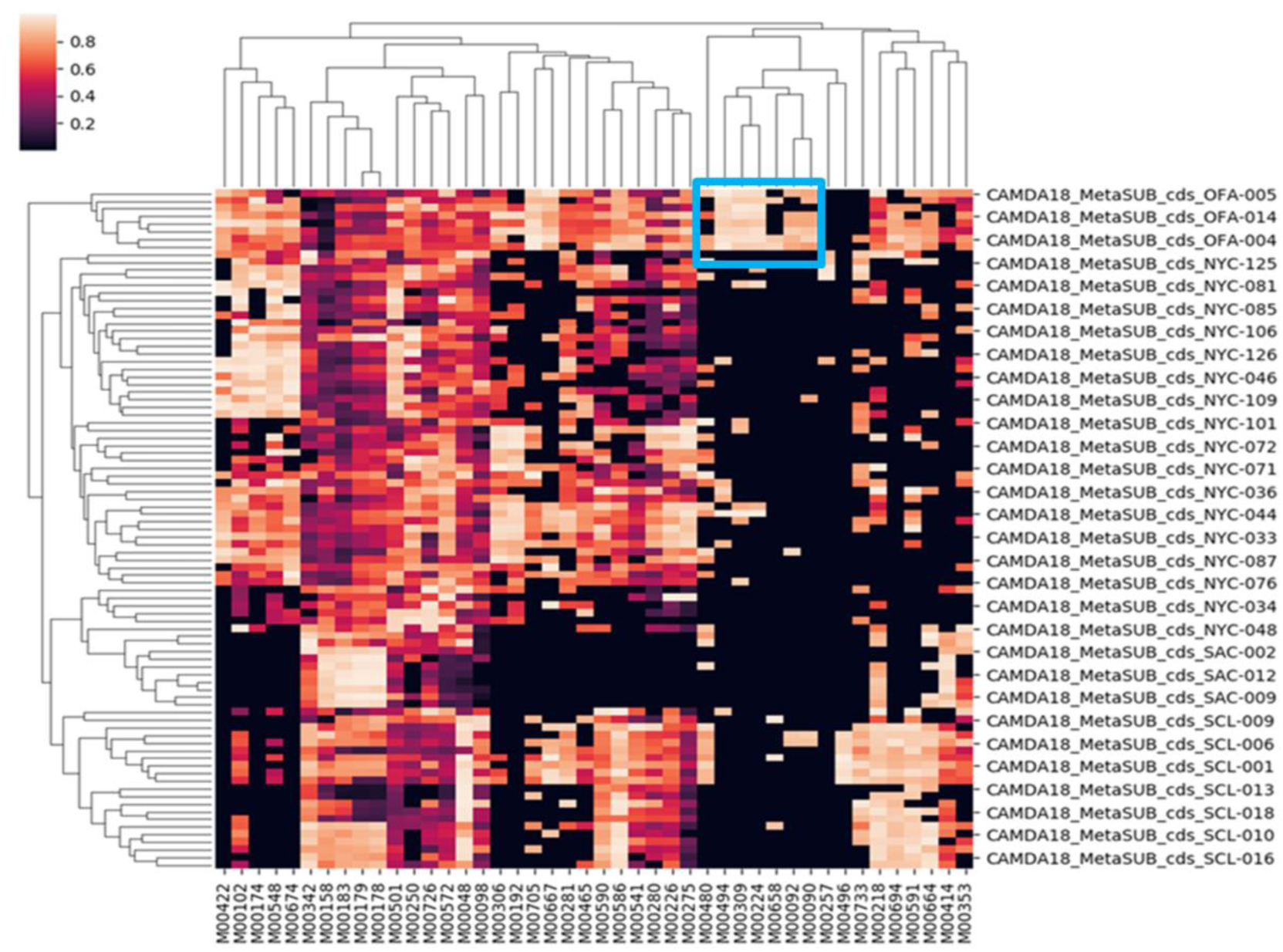
The most relevant KEGG features extracted from the classification pipeline by averaging the feature importance of each base learner of the ensemble in each run of the experiment. In a blue square the features characteristic from Ofa, and listed in table 2, are shown.

Particularly, *Fluoroquinolone transport system* (M00224) is an ABC-2 type transporter that confers resistance to fluoroquinolone, a widely used antibiotic [31, 32]. Similarly, *VraS-VraR* (M00480) and *VanS-VanR* (M00658) are two-component regulatory systems involved in the response to two antibiotics, β-lactam [33] and glycopeptides [34], respectively. Interestingly, *Fluoroquinolone transport system* and *VraS-VraR* are known to confer resistance in *Staphylococcus aureus*, a pathogen which is known to have higher incidence rates in sub Saharan Africa than those reported from developed countries [35], maybe because a higher genetic susceptibility of these populations [36]. Since *Staphylococcus aureus* is a skin pathogen it is easier to find it over-represented in the African MetaSUB samples. This observation captured by the functional analysis of MetaSUB samples proposed here suggests an excessive use of antibiotics that could eventually have caused an emergence of resistant strains. Actually, epidemiologic studies report the prevalence of Staphylococcal disease in sub-Saharan Africa, along with an increase in antibiotic resistance [35]. Moreover, two single-nucleotide polymorphisms (SNPs) in the human leukocyte antigen (HLA) class II region on chromosome 6 was demonstrated to be associated with susceptibility to *S. aureus* infection at a genome-wide significant level [37] and a recent admixture mapping study demonstrated that genomic variations with different frequencies in these SNPs in European and African ancestral genomes influence susceptibility to *S. aureus* infection, strongly suggesting a genetic basis for our observations [36].

### Classification of new samples of the cities in the training set

In order to test the generalization power of the predictor obtained with the *training dataset*, we have used the *test dataset 1* composed by 30 samples belonging to the same cities that the *training dataset*. Table 3 shows the cross validation and the confusion matrix, in which, the functional heterogeneity of New York clearly introduces some noise in the classification (probably with a real biological meaning). The accuracy of the predictor is of 0.73.

**Table 3.** Cross validation and confusion matrix of KEGG functional profiles obtained from the samples from the test dataset 1, belonging to the cities from the training dataset.

| Truth / Preds | Auckland | Hamilton | NY | Ofa | Porto | Sacramento | Santiago | Tokyo | All | Accuracy |
| --- | --- | --- | --- | --- | --- | --- | --- | --- | --- | --- |
| NY | 1 | 1 | 8 | 0 | 0 | 0 | 0 | 0 | 10 | 0.8 |
| Ofa | 0 | 0 | 2 | 3 | 0 | 0 | 0 | 0 | 5 | 0.6 |
| Porto | 0 | 0 | 1 | 0 | 8 | 0 | 0 | 1 | 10 | 0.8 |
| Santiago | 0 | 0 | 1 | 0 | 0 | 1 | 3 | 0 | 5 | 0.6 |
| All | 1 | 1 | 12 | 3 | 8 | 1 | 3 | 1 | 30 | 0.73 |

### Classification using different functional profiles

KEGG encompasses a global compendium of bacterial functionalities, providing features with a high discriminatory power but, in some cases, with no much biological interest, which can mask functionalities of more relevance from a medical, forensic or epidemiological viewpoint. Instead, other databases that collect specific bacterial activities or functionalities could be used. Since antibiotic resistance has emerged among the generic functionalities as a high relevant feature in the classification, in addition to have an obvious importance by itself, it seemed worth focusing on features that specifically describe antibiotic resistances. Therefore, a new training process was carried out using CARD, the database of antibiotic resistances [24]. Again, a set of antibiotic resistance features clearly distinguishes Ofa from the rest of cities, as previously observed (Figure 5A). Table 4 describes the specific resistances found as characteristic from Ofa which, overall, reinforce our previous finding with KEGG about transporters [31, 32] and two-component regulatory systems involved in the response to antibiotics [33, 34], but providing more detail on specific resistance mechanisms. Interestingly, the characteristic that distinguishes Porto samples from those of other cities is the absence of antibiotic resistances (Figure 5B). Although we do not have a strong epidemiological explanation for this, recent studies show that Portugal is among the countries in Europe with the highest defined daily dose per habitant [38]. Whether the high antibiotic consumption can be behind this observation or not needs of deeper epidemiological studies but, in any case, it is pointing to a distinctive local characteristic of clear epidemiological relevance.

**Table 4.** The most relevant antibiotic resistance modules (CARD) in Ofa

| ACCESSION | NAME | DESCRIPTION |
| --- | --- | --- |
| 3002940 | vanSN | vanSN is a vanS variant found in the vanN gene cluster |
| 3000217 | blaR1 | blaR1 is a transmembrane spanning and signal transducing protein which in response to interaction with beta-lactam antibiotics results in upregulation of the blaZ/blaR1/blaI operon. |
| 3003069 | vanXYG | vanXYG is a vanXY variant found in the vanG gene cluster |
| 3000180 | tetA(P) | TetA(P) is a inner membrane tetracycline efflux protein found on the same operon as the ribosomal protection protein TetB(P). It is found in Clostridium, a Gram-positive bacterium. |
| 3002541 | AAC(3)-VIIa | AAC(3)-VIIa is a chromosomal-encoded aminoglycoside acetyltransferase in Streptomyces rimosus |

**Figure 5.**
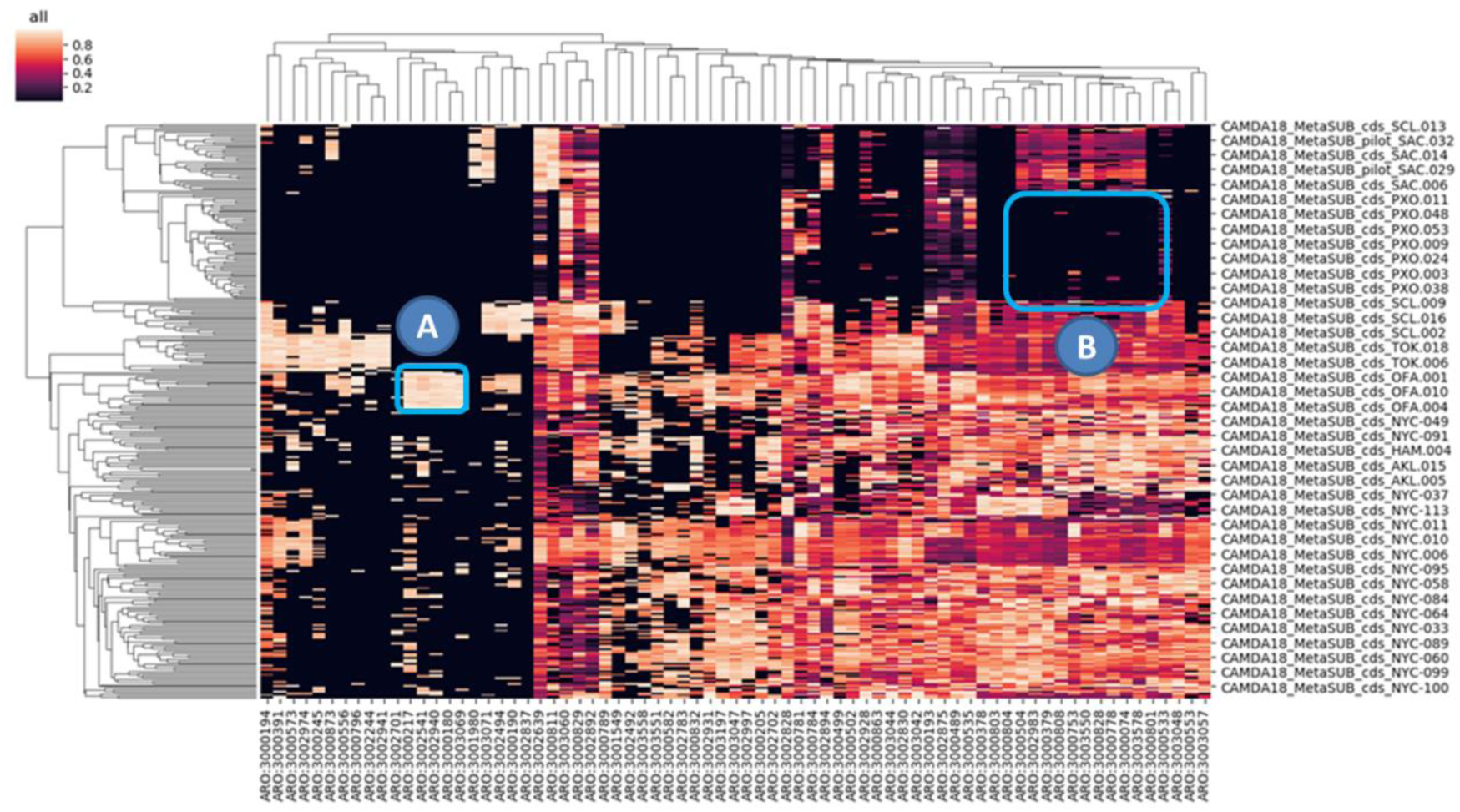
The most relevant CARD (antibiotic resistances) features extracted from the classification pipeline by averaging the feature importance of each base learner of the ensemble in each run of the experiment. A) Features characteristic from Ofa. B) Features characteristic from Porto.

Table 5 shows the cross validation and the confusion matrix with the CARD functional profiles, in which, the functional heterogeneity of New York is still introducing some noise in the classification but the accuracy of the predictor increased to 0.8.

**Table 5.** Cross validation and confusion matrix of antibiotic resistances (CARD) functional profiles obtained from the samples from the test dataset 1, belonging to the cities from the training dataset.

| Truth/pred | Auckland | NY | Ofa | Porto | Santiago | All | Accuracy |
| --- | --- | --- | --- | --- | --- | --- | --- |
| NY | 2 | 8 | 0 | 0 | 0 | 10 | 0.8 |
| Ofa | 0 | 1 | 4 | 0 | 0 | 5 | 0.8 |
| Porto | 0 | 0 | 0 | 10 | 0 | 10 | 1 |
| Santiago | 0 | 2 | 0 | 1 | 2 | 5 | 0.4 |
| All | 2 | 11 | 4 | 11 | 2 | 30 | 0.8 |

### Classification using mixed functional profiles

In addition to build predictors with a single functional feature, it is possible to combine different functional profiles producing higher accuracies in the classification. Here we combined KEGG and CARD profiles using the Fusion Pipeline (see Methods) and the resulting classification accuracy increased to 0.9. Table 6 shows the cross validation values obtained with the mixed profiles. Only New York, which is the most heterogeneous cite from a functional point of view, shows a couple of bad predictions (the Ofa misplaced sample was assigned to New York, probably for the same reason).

**Table 6.** Cross validation and confusion matrix of functional profiles obtained from the combination of KEGG and CARD corresponding to samples from the test dataset 1 belonging to the cities from the training dataset.

| truth/pred | Auckland | NY | Ofa | Porto | Santiago | All | Accuracy |
| --- | --- | --- | --- | --- | --- | --- | --- |
| NY | 1 | 8 | 1 | 0 | 0 | 10 | 0.8 |
| Ofa | 0 | 1 | 4 | 0 | 0 | 5 | 0.8 |
| Porto | 0 | 0 | 0 | 10 | 0 | 10 | 1 |
| Santiago | 0 | 0 | 0 | 0 | 5 | 5 | 1 |
| All | 1 | 11 | 3 | 10 | 5 | 30 | 0.9 |

More functional profiles could be included by using an extension of the Fusion Pipeline to N datasets as previously shown [39], coupled with robust Least Squares techniques [40], to accommodate for the challenging low sample size high dimensional data scenario.

### Classification new samples of with new cities

In order to check the performance of the predictor with samples from cities that were not used in the initial *training dataset* we used the 30 samples from the *test dataset 2*, from the cities: Ilorin (close to Ofa), Lisbon (in Portugal as Porto, but not close) and Boston (in USA, but not close to Ney York).

Figure 6 shows how the cities are clustered very much as expected. Thus Ilorin maps together with Ofa (Figure 6A) because these two cities are physically close cities in Nigeria (and connected by a train). As expected, the New York cluster shows the highest dispersion (Figure 6B), although is not similar to Boston (Figure 6C). The same is observed with Lisbon (Figure 6D), which is not close to Porto (Figure 6E) and both map in different places. Interestingly, the Porto “outlier” sample maps on the Lisbon cluster. Similarly to the case of Ofa and Ilorin, Hamilton and Auckland, both New Zealand cities connected by a train also map together as well (Figure 6F).

**Figure 6.**
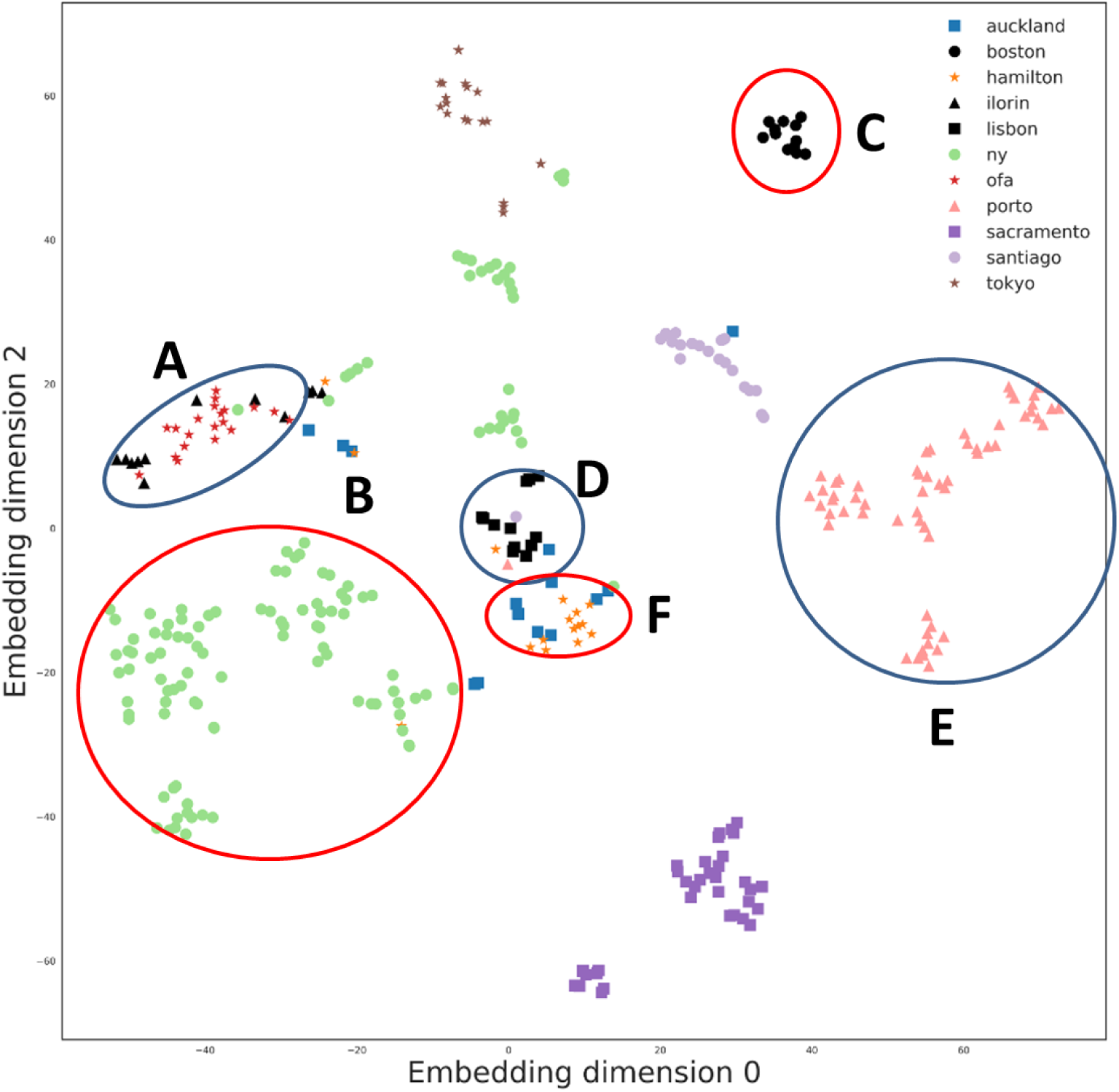
Classification of all the cities based on KEGG-based functional profiles. A) Ilorin and Ofa, two physically close cities in Nigeria (connected by a train) map close to each other. B) New York, not close to C) Boston, D) Lisbon is not close to E) Porto. F) Hamilton and Auckland, both New Zealand cities connected by a train, also map together.

### Machine Learning Pipeline Comparison

Finally, the performance of each machine learning pipeline was evaluated by joining the samples from the training and the three validation datasets. For each model a 10-fold city-wise stratified cross-validation was performed. In order to provide statistical evidence for the results each experiment is repeated 10 times with different random seeds initializations. Figure 7 shows a box plot diagram of the different experiments grouped by the functional profile used, namely: *kegg* for KEGG-Modules, *card* for CARD-ARO and *fusion* for the Multiview case. As expected, the model performance follows the tendency already exhibited: the fusion pipeline outperforms the single-view case, and the CARD-ARO *view* provides slightly better results than KEGG-Modules.

**Figure 7.**
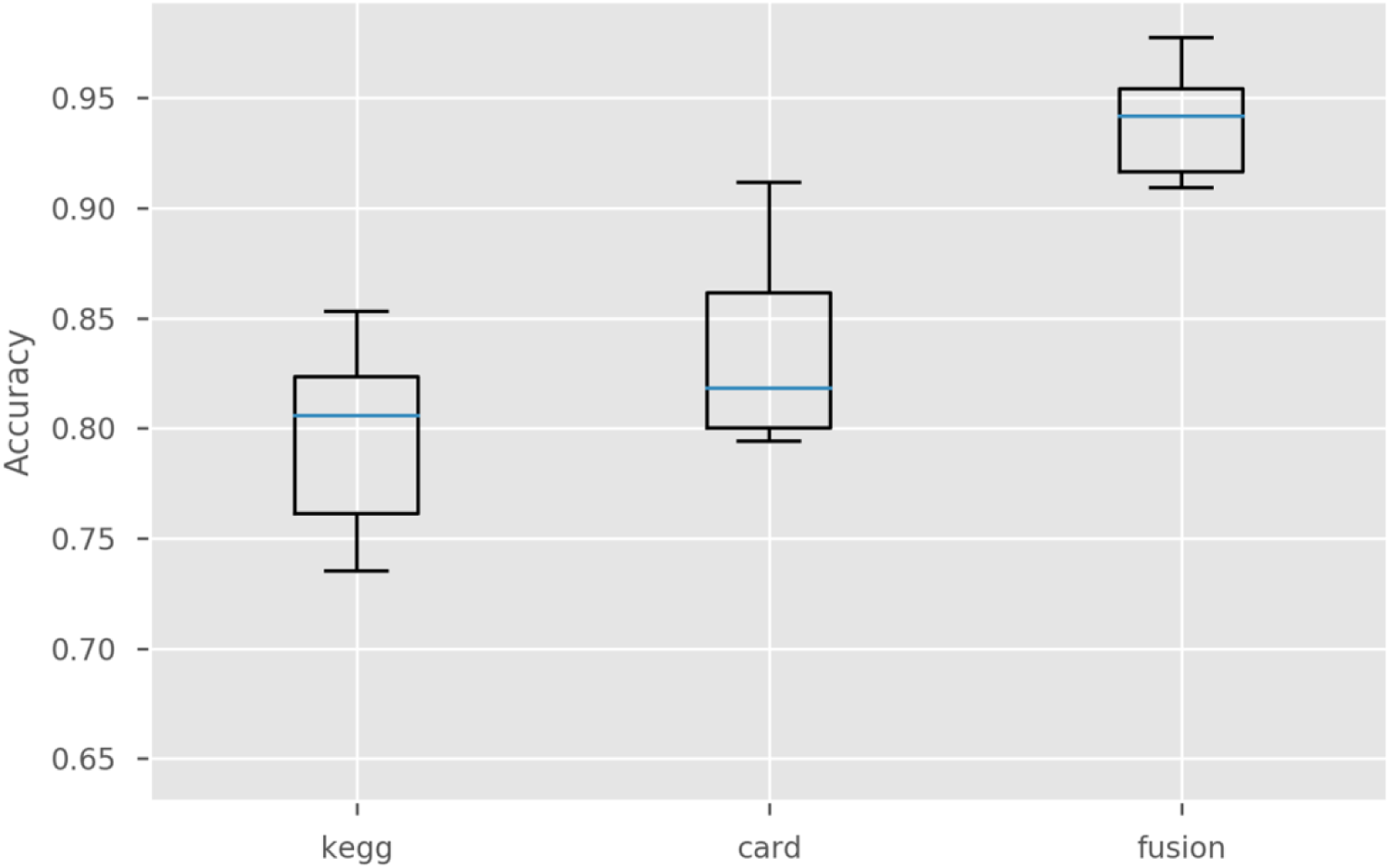
Accuracies obtained using the whole dataset (Training dataset and test datasets 1, 2 and 3) with only KEGG profiles, only CARD profiles and the fusion of both profiles.

## Conclusions

The recodification of metagenomics data from the conventional gene or strain abundance profiles to other types of profiles with biological meaning offers new avenues for the analysis of microbiome data. Here we show how the use of KEGG- and CARD-based functional profiles, derived from the original metagenomics data, not only provides accurate sample classification but also offers interesting epidemiological and biological interpretations of the results found. Interestingly, antibiotic resistance arises as a relevant classification feature, supported by epidemiological [35] and genetic [36] previous observations.

## List of abbreviations

CAMDA: Critical Assessment of Massive Data Analysis
CARD: Comprehensive Antibiotic Resistance Database
CCA: Canonical Correlation Analysis
HLA: Human Leukocyte Antigen
KEGG: Kyoto Encyclopedia of Genes and Genomes
PCA: Principal Component Analysis
SNP: Single Nucleotide Polymorphisms
t-SNE: t-distributed Stochastic Neighbor Embedding
WGS: whole genome sequencing

## Declarations

### Ethical Approval and Consent to participate

Not applicable

### Consent for publication

Not applicable

### Availability of supporting data

Data sharing is not applicable to this article as no datasets were generated during the current study

## Author’s contributions

CSCS carried out the data preprocessing and interpretation parts of the analysis; CL performed the machine learning part of the analysis; JPF and DLL helped in different parts of the analysis of the results; JD conceived the work and wrote the paper. All authors read and approved the final manuscript

## Acknowledgements

We appreciate very much the advices of Dr. Jaime Huerta-Cepas regarding data processing.

## References

1. Cho I, Blaser MJ: The human microbiome: at the interface of health and disease. Nature Reviews Genetics 2012, 13(4):260.

2. Garrido-Cardenas JA, Manzano-Agugliaro F: The metagenomics worldwide research. Current genetics 2017, 63(5):819–829.

3. Gilbert JA, Jansson JK, Knight R: The Earth Microbiome project: successes and aspirations. BMC biology 2014, 12(1):69.

4. Mason C, Afshinnekoo E, Ahsannudin S, Ghedin E, Read T, Fraser C, Dudley J, Hernandez M, Bowler C, Stolovitzky G: The Metagenomics and Metadesign of the Subways and Urban Biomes (MetaSUB) International Consortium inaugural meeting report. MICROBIOME 2016, 4(1):24.

5. Snel B, Bork P, Huynen MA: Genome phylogeny based on gene content. Nature genetics 1999, 21(1):108.

6. Zaneveld JR, Lozupone C, Gordon JI, Knight R: Ribosomal RNA diversity predicts genome diversity in gut bacteria and their relatives. Nucleic acids research 2010, 38(12):3869–3879.

7. Langille MG, Zaneveld J, Caporaso JG, McDonald D, Knights D, Reyes JA, Clemente JC, Burkepile DE, Thurber RLV, Knight R: Predictive functional profiling of microbial communities using 16S rRNA marker gene sequences. Nature biotechnology 2013, 31(9):814.

8. Tringe SG, Hugenholtz P: A renaissance for the pioneering 16S rRNA gene. Current opinion in microbiology 2008, 11(5):442–446.

9. Segata N, Boernigen D, Tickle TL, Morgan XC, Garrett WS, Huttenhower C: Computational meta'omics for microbial community studies. Molecular systems biology 2013, 9(1):666.

10. Tyson GW, Chapman J, Hugenholtz P, Allen EE, Ram RJ, Richardson PM, Solovyev VV, Rubin EM, Rokhsar DS, Banfield JF: Community structure and metabolism through reconstruction of microbial genomes from the environment. Nature 2004, 428(6978):37.

11. Quince C, Walker AW, Simpson JT, Loman NJ, Segata N: Shotgun metagenomics, from sampling to analysis. Nature biotechnology 2017, 35(9):833.

12. Scholz M, Ward DV, Pasolli E, Tolio T, Zolfo M, Asnicar F, Truong DT, Tett A, Morrow AL, Segata N: Strain-level microbial epidemiology and population genomics from shotgun metagenomics. Nature methods 2016, 13(5):435.

13. Börnigen D, Morgan XC, Franzosa EA, Ren B, Xavier RJ, Garrett WS, Huttenhower C: Functional profiling of the gut microbiome in disease-associated inflammation. Genome medicine 2013, 5(7):65.

14. Lloyd-Price J, Mahurkar A, Rahnavard G, Crabtree J, Orvis J, Hall AB, Brady A, Creasy HH, McCracken C, Giglio MG: Strains, functions and dynamics in the expanded Human Microbiome Project. Nature 2017, 550(7674):61.

15. Zolfo M, Asnicar F, Manghi P, Pasolli E, Tett A, Segata N: Profiling microbial strains in urban environments using metagenomic sequencing data. Biology direct 2018, 13(1):9.

16. Kultima JR, Coelho LP, Forslund K, Huerta-Cepas J, Li SS, Driessen M, Voigt AY, Zeller G, Sunagawa S, Bork P: MOCAT2: a metagenomic assembly, annotation and profiling framework. Bioinformatics 2016, 32(16):2520–2523.

17. Cox MP, Peterson DA, Biggs PJ: SolexaQA: At-a-glance quality assessment of Illumina second-generation sequencing data. BMC Bioinformatics 2010, 11(1):485.

18. Li R, Yu C, Li Y, Lam T-W, Yiu S-M, Kristiansen K, Wang J: SOAP2: an improved ultrafast tool for short read alignment. Bioinformatics 2009, 25(15):1966–1967.

19. Hyatt D, Chen G-L, LoCascio PF, Land ML, Larimer FW, Hauser LJ: Prodigal: prokaryotic gene recognition and translation initiation site identification. BMC Bioinformatics 2010, 11(1):119.

20. Fu L, Niu B, Zhu Z, Wu S, Li W: CD-HIT: accelerated for clustering the next-generation sequencing data. Bioinformatics 2012, 28(23):3150–3152.

21. Buchfink B, Xie C, Huson DH: Fast and sensitive protein alignment using DIAMOND. Nature methods 2015, 12(1):59.

22. Huerta-Cepas J, Szklarczyk D, Forslund K, Cook H, Heller D, Walter MC, Rattei T, Mende DR, Sunagawa S, Kuhn M: eggNOG 4.5: a hierarchical orthology framework with improved functional annotations for eukaryotic, prokaryotic and viral sequences. Nucleic acids research 2015, 44(D1):D286–D293.

23. Kanehisa M, Goto S, Sato Y, Kawashima M, Furumichi M, Tanabe M: Data, information, knowledge and principle: back to metabolism in KEGG. Nucleic Acids Res 2014, 42(Database issue):D199–205.

24. Jia B, Raphenya AR, Alcock B, Waglechner N, Guo P, Tsang KK, Lago BA, Dave BM, Pereira S, Sharma AN: CARD 2017: expansion and model-centric curation of the comprehensive antibiotic resistance database. Nucleic acids research 2016:gkw1004.

25. Pedregosa F, Varoquaux G, Gramfort A, Michel V, Thirion B, Grisel O, Blondel M, Prettenhofer P, Weiss R, Dubourg V: Scikit-learn: Machine learning in Python. Journal of machine learning research 2011, 12(Oct):2825–2830.

26. Maaten Lvd, Hinton G: Visualizing data using t-SNE. Journal of machine learning research 2008, 9(Nov):2579–2605.

27. Chen T, Guestrin C: Xgboost: A scalable tree boosting system. In: Proceedings of the 22nd acm sigkdd international conference on knowledge discovery and data mining: 2016. ACM: 785–794.

28. Snoek J, Larochelle H, Adams RP: Practical bayesian optimization of machine learning algorithms. In: Advances in neural information processing systems: 2012. 2951–2959.

29. Sun Q-S, Zeng S-G, Liu Y, Heng P-A, Xia D-S: A new method of feature fusion and its application in image recognition. Pattern Recognition 2005, 38(12):2437–2448.

30. Papadopoulos CJ, Carson CF, Chang BJ, Riley TV: Role of the MexAB-OprM efflux pump of Pseudomonas aeruginosa in tolerance to tea tree (Melaleuca alternifolia) oil and its monoterpene components terpinen-4-ol, 1, 8–cineole, and α-terpineol. Applied and environmental microbiology 2008, 74(6):1932–1935.

31. Hooper DC: Emerging mechanisms of fluoroquinolone resistance. Emerging infectious diseases 2001, 7(2):337.

32. Kaatz GW, Seo SM, Ruble CA: Efflux-mediated fluoroquinolone resistance in Staphylococcus aureus. Antimicrobial agents and chemotherapy 1993, 37(5):1086–1094.

33. Boyle-Vavra S, Yin S, Daum RS: The VraS/VraR two-component regulatory system required for oxacillin resistance in community-acquired methicillin-resistant Staphylococcus aureus. FEMS microbiology letters 2006, 262(2):163–171.

34. Arthur M, Molinas C, Courvalin P: The VanS-VanR two-component regulatory system controls synthesis of depsipeptide peptidoglycan precursors in Enterococcus faecium BM4147. Journal of bacteriology 1992, 174(8):2582–2591.

35. Herrmann M, Abdullah S, Alabi A, Alonso P, Friedrich AW, Fuhr G, Germann A, Kern WV, Kremsner PG, Mandomando I: Staphylococcal disease in Africa: another neglected ‘tropical’disease. Future microbiology 2013, 8(1):17–26.

36. Cyr D, Allen A, Du G, Ruffin F, Adams C, Thaden J, Maskarinec S, Souli M, Guo S, Dykxhoorn D: Evaluating genetic susceptibility to Staphylococcus aureus bacteremia in African Americans using admixture mapping. Genes and immunity 2017, 18(2):95.

37. DeLorenze GN, Nelson CL, Scott WK, Allen AS, Ray GT, Tsai A-L, Quesenberry Jr CP, Fowler Jr VG: Polymorphisms in HLA class II genes are associated with susceptibility to Staphylococcus aureus infection in a white population. The Journal of infectious diseases 2015, 213(5):816–823.

38. Goossens H, Ferech M, Vander Stichele R, Elseviers M, Group EP: Outpatient antibiotic use in Europe and association with resistance: a cross-national database study. The Lancet 2005, 365(9459):579–587.

39. Kettenring JR: Canonical analysis of several sets of variables. Biometrika 1971, 58(3):433–451.

40. Bilenko NY, Gallant JL: Pyrcca: regularized kernel canonical correlation analysis in python and its applications to neuroimaging. Frontiers in neuroinformatics 2016, 10:49.

